# CryoARC: Atomic-resolution conformational landscapes of protein assemblies from cryo-EM single particles with evolutionary priors

**DOI:** 10.64898/2026.05.25.727696

**Authors:** Rémi Vuillemot, Sergei Grudinin

## Abstract

Single-particle cryo-electron microscopy (cryo-EM) reveals structural heterogeneity in macromolecular complexes, but recovering continuous conformational landscapes at high resolution remains challenging. Here, we introduce CryoARC, a deep learning framework that integrates evolutionary sequence representations with cryo-EM particle images to reconstruct continuous conformational ensembles at atomic resolution. CryoARC combines a latent representation of particle heterogeneity with a sequence-conditioned structure decoder inspired by protein structure prediction architectures, enabling direct prediction of particle-specific atomic structures. We further introduce a heterogeneous refinement strategy that aggregates per-particle predictions into a canonical density map, improving reconstruction quality and resolution. We evaluate CryoARC on both synthetic and experimental datasets and show that it recovers continuous conformational landscapes together with coherent atomic models. CryoARC demonstrates how sequence-derived structural priors can be combined with cryo-EM particle images for ensemble-based atomic reconstruction of heterogeneous macromolecular systems. CryoARC is fully open source and available at https://gricad-gitlab.univ-grenoble-alpes.fr/GruLab/CryoARC.

## Introduction

Cryo-electron microscopy (cryo-EM) allows the study of biological macromolecules in near-native conditions, where multiple conformations coexist in solution. Each particle image corresponds to a noisy projection of a single molecular state, and together these images define an underlying conformational landscape.

Recent advances in cryo-EM data analysis have improved the characterization of structural heterogeneity. Methods have progressed from discrete classification-based reconstructions [1, 2] to continuous representations of conformational variability [3–18]. These approaches learn low-dimensional latent representations of the noisy particle images and map them into three-dimensional structures. However, the low signal-to-noise ratio of cryo-EM images makes it difficult to disentangle conformational variability from imaging noise. Consequently, many methods incorporate prior information through regularization, including physics-inspired and structure-based constraints [6–8, 16–18].

In parallel, recent advances in protein structure prediction have demonstrated that evolutionary sequence data encodes rich structural information [19–22]. Sequence representations derived from multiple sequence alignments and protein language models are effective not only in capturing structural organization, but also in representing conformational variability [23–35]. Despite their success in structure prediction, the integration of sequence-derived priors into cryo-EM heterogeneity reconstruction remains largely unexplored.

In CryoARC, we incorporate evolutionary sequence information into cryo-EM heterogeneity reconstruction within a deep learning framework. We use embeddings derived from multiple sequence alignments as sequence-based priors that encode evolutionary constraints related to conformational variability. These embeddings are integrated into an encoder–decoder architecture that maps particle images to a latent representation and decodes each particle into an atomic structure. As a result, CryoARC represents continuous conformational landscapes at atomic resolution, without requiring an input structure or explicit physics-based models. We further introduce a heterogeneous refinement strategy that aggregates particle-specific atomic predictions into a canonical high-resolution density map. This refinement improves the quality of reconstruction across heterogeneous states compared to conventional homogeneous reconstruction approaches.

We evaluated CryoARC on synthetic and experimental datasets, including a public benchmark for continuous heterogeneity methods and two EMPIAR datasets that span molecular weights from 102 to 164 kDa. CryoARC recovers continuous conformational landscapes and produces coherent particle-specific atomic models across heterogeneous states. It shows improved reconstruction quality compared to recent continuous heterogeneity methods. While being currently limited to protein complexes up to a few thousand residues due to computational scaling, CryoARC enables particle-specific atomic modeling of continuous conformational variability by integrating sequence-informed priors with cryo-EM data.

### Highlights

- CryoARC incorporates evolutionary sequence information from multiple sequence alignments into cryo-EM heterogeneity reconstruction.
- CryoARC predicts atomic-level continuous conformational landscapes from cryo-EM images and protein sequences.
- A heterogeneous refinement strategy aggregates particle-level atomic predictions into consensus density maps.
- The approach is demonstrated on synthetic and experimental datasets.

## Results

### Overview

CryoARC models continuous conformational variability from single-particle cryo-EM datasets by integrating evolutionary sequence information into the image-based reconstruction process. Given a set of cryo-EM particle images 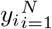 and the aminoacid sequence of the macromolecular complex, we train an end-to-end differentiable model that infers a low-dimensional latent variable *z*_*i*_ describing the conformational state of each particle from the images. An all-atom structure *s*_*i*_ is then reconstructed by a decoder conditioned on both the latent variable and sequence-derived evolutionary priors.

The architecture consists of three components. A sequence encoder processes multiple sequence alignments (MSAs) using a frozen pretrained Evoformer, producing residue-wise and pairwise embeddings (*h, p*) that encode evolutionary structural information. An image encoder maps each particle image to a probabilistic latent variable *z*_*i*_, capturing conformational variability in the dataset. A structure decoder then generates atomic structures *s*_*i*_ by conditioning sequence-derived embeddings on the latent variable through an attention-based architecture. Figure 1 provides an overview of the CryoARC architecture.

**Fig. 1.**
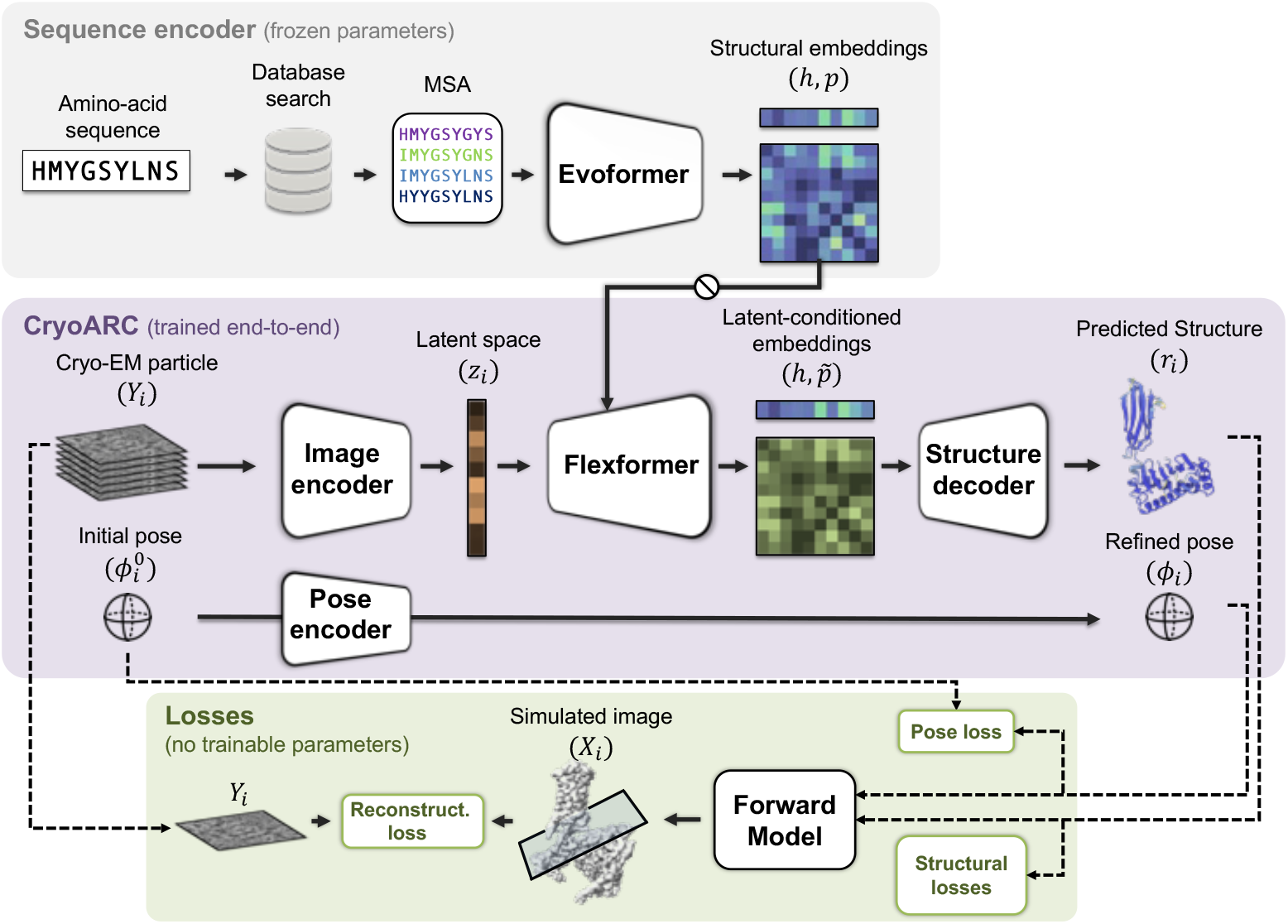
Schematic workflow of the CryoARC model. The sequence encoder processes the amino-acid sequence through multiple sequence alignment (MSA) and a pretrained Evoformer network to generate structural embeddings. The CryoARC model encodes cryo-EM particle images to infer a latent representation of conformational variability, which is then combined with the sequence-derived embeddings in a structure decoder to generate atomic models. The bottom panel illustrates the training objectives, including reconstruction loss, pose estimation loss, and structural regularization terms.

During training, CryoARC converts predicted atomic structures into cryo-EM particle images using a differentiable forward model of image formation that simulates microscope acquisition. The simulated images are compared with the observed particle images through the reconstruction loss. The entire pipeline is trained end-to-end in a variational framework, with the exception of the sequence encoder, which is pretrained on a large corpus of protein structures and kept frozen during training. Since sequence embeddings are independent of individual particle images, they are computed once and reused across all particles.

After training, the predicted particle-specific structures enable downstream analysis of conformational variability. In particular, the latent representations can be used to reconstruct continuous trajectories that describe transitions between conformational states, offering insight into functional dynamics. We also introduce a heterogeneous refinement strategy that constructs a consensus density map from particle-specific atomic predictions. Each particle contribution is placed into a common reference frame defined by its predicted structure and aggregated to produce a refined density map.

### Understanding conformational variability helps increasing resolution

For the demonstration of our approach, we selected challenging datasets with conformational heterogeneity of protein complexes in the range of 102 to 164 kDa. We first applied the method to the HER2–trastuzumab–pertuzumab complex (EMPIAR-11665, Fig. 2a) [36], a 164 kDa antibody–antigen assembly exhibiting pronounced conformational flexibility in the antibody regions. Previous analyses of this dataset have demonstrated that discrete classification methods are insufficient to fully resolve its structural variability and that continuous approaches are required to describe its conformational landscape [36]. In particular, the complex is characterized by a relatively rigid HER2–pertuzumab core and a flexible trastuzumab region. Previous studies identified dominant modes of motion corresponding to hinge-like rotations of trastuzumab relative to the core, as well as an additional axial rotation.

**Fig. 2.**
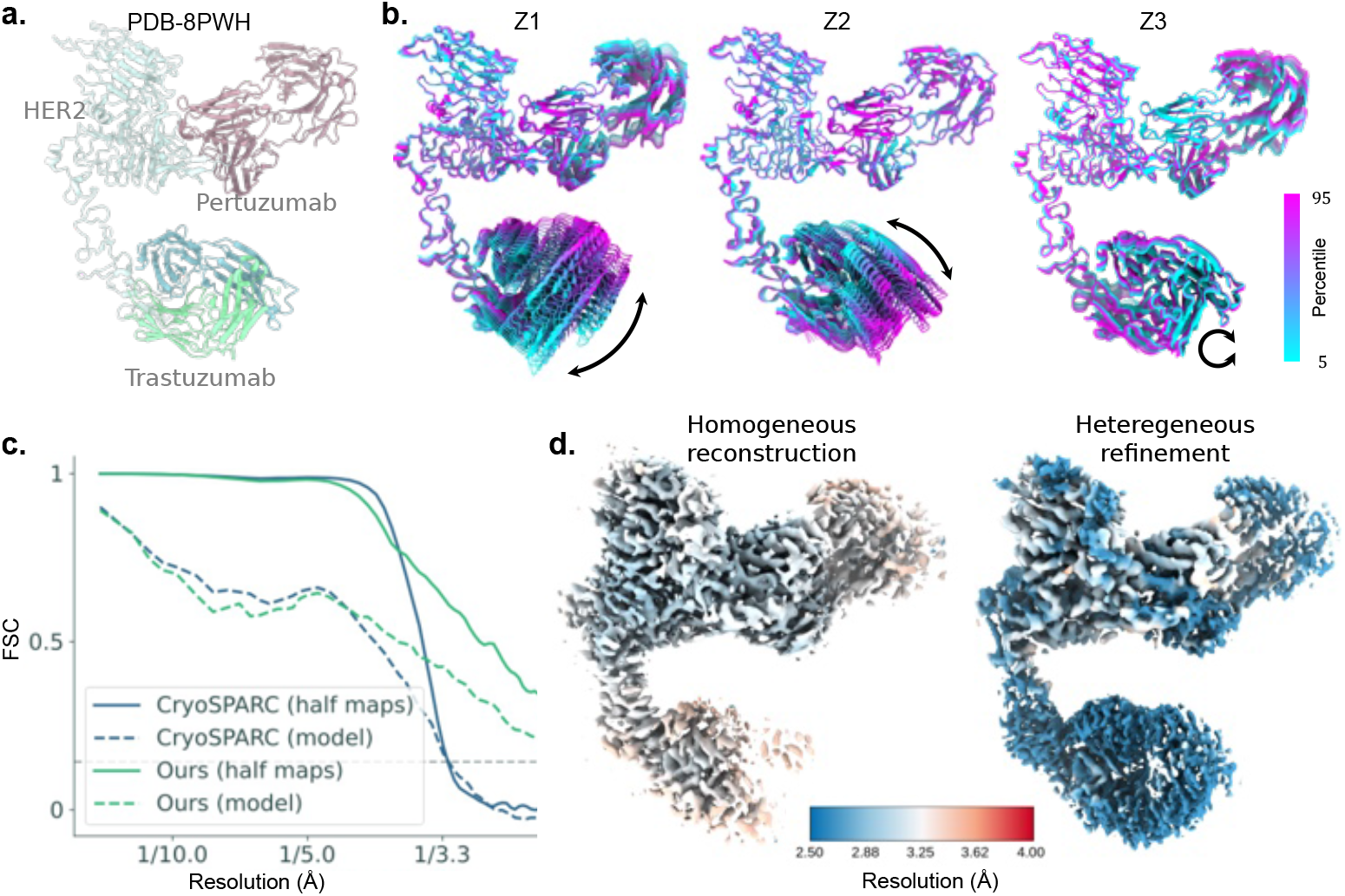
CryoARC results on the HER2 dataset. **a**. Density map and corresponding atomic model of the HER2–trastuzumab–pertuzumab complex. **b**. Conformational trajectories along the first three latent dimensions (Z1–Z3), obtained by varying each latent variable between the 5th and 95th percentiles while keeping the others fixed. **c**. Fourier shell correlation (FSC) curves comparing homogeneous reconstruction and heterogeneous refinement, including half-map FSC and model-to-map FSC. **d**. Density maps for homogeneous reconstruction and heterogeneous refinement, colored by local resolution estimated from FSC.

Figure 2 reports CryoARC results on EMPIAR-11665. The analysis of the learned conformational space (Fig. 2b) recovers the three dominant modes of motion of the trastuzumab region (Z1–Z3). Z1 and Z2 describe hinge-like rotations of trastuzumab relative to the HER2–pertuzumab core, while Z3 captures a rotation around its own axis. The recovered conformational variability is consistent with the motions reported in the original study.

Using the inferred heterogeneity, we apply the heterogeneous refinement strategy described above. The resulting reconstruction outperforms the homogeneous reconstruction, as demonstrated by Fourier Shell Correlation (FSC) analysis (Fig. 2c), indicating an improvement in global resolution. Local resolution maps further show improved density in the flexible trastuzumab region (Fig. 2d). A comparative analysis of HER2 datasets (Fig. S1) against homogeneous reconstruction (EMD-18188) and multibody refinement (EMD-17993) reveals greater clarity in flexible regions, including more distinct side-chain density in CryoARC reconstructions.

### CryoARC reconstructs atomic models of missing regions

We next applied CryoARC to an experimental cryo-EM dataset (EMPIAR-10330) consisting of particles of the Plasmodium falciparum chloroquine resistance transporter (PfCRT) 7G8 isoform in complex with a Fab fragment [37]. PfCRT is a 49 kDa membrane transporter involved in antimalarial drug resistance, and the Fab-bound construct increases the overall size and stability of the complex for cryo-EM analysis. The original reconstruction used 16,905 particles to obtain a 3.2 Å map (EMD-20806) and an associated atomic model (PDB-6UKJ), in which the Fab constant domain is not resolved due to its conformational flexibility (Fig. 3a). The original structure reveals a transmembrane core with an extracellular Fab-binding region, containing only the Fab variable domain.

**Fig. 3.**
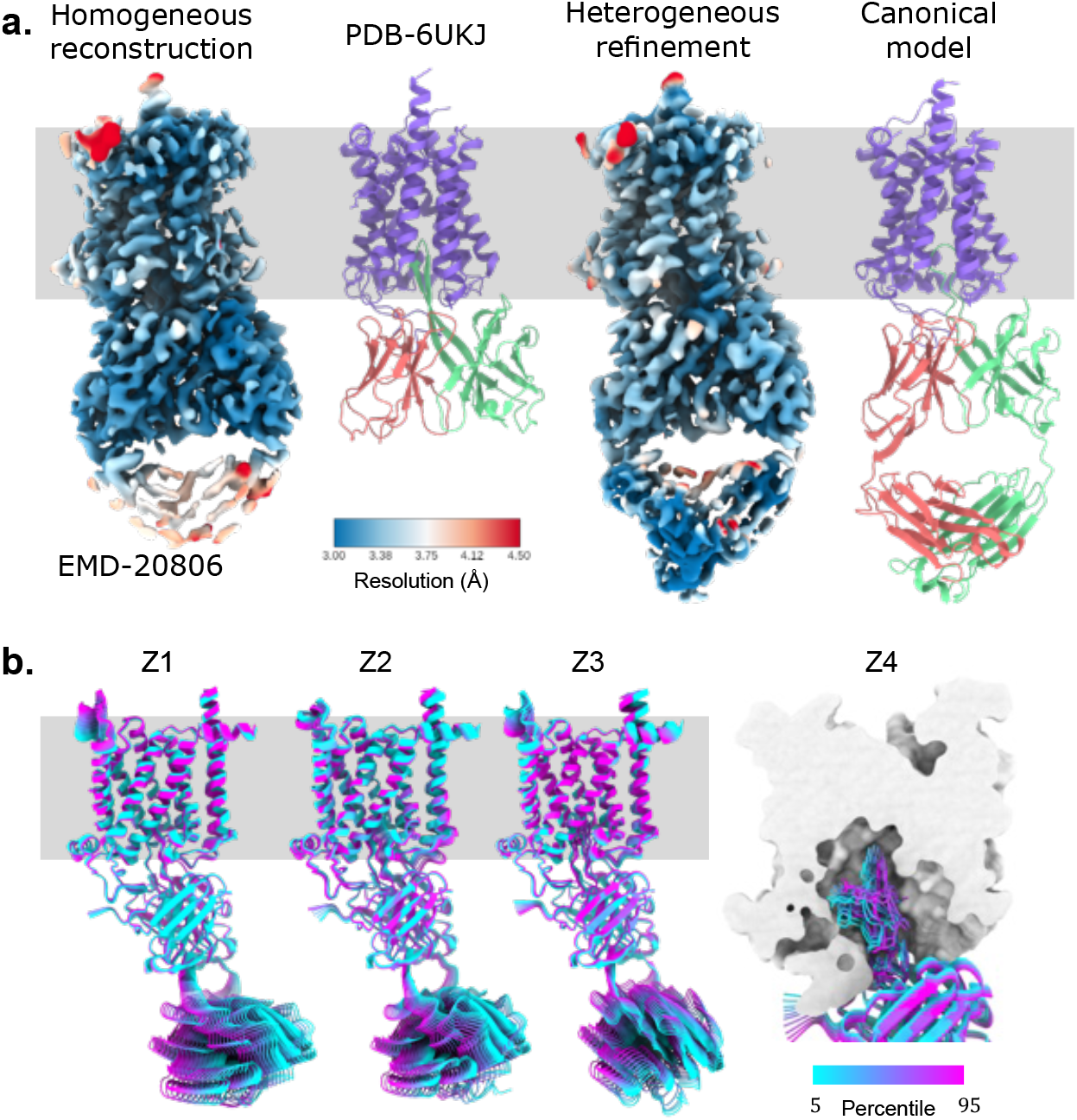
CryoARC results on the PfCRT dataset. **a**. Density map and the corresponding atomic model for the homogeneous reconstruction (EMD-20806; PDB-6UKJ, left). The heterogeneous refinement result is shown on the right, with the reconstructed map colored by local resolution estimated from Fourier shell correlation (FSC), and the corresponding canonical model. **b**. Conformational trajectories along the first four latent dimensions (Z1–Z4). Z1–Z3 primarily describe motions of the Fab constant domain, while Z4 captures variability in the Fab variable-region loop; only the Fab region is shown as ribbons for Z4 for clarity. Trajectories are generated by varying each latent dimension between the 5th and 95th percentiles while keeping others fixed.

We use this dataset to evaluate CryoARC on the reconstruction of conformationally heterogeneous regions, focusing particularly on the unresolved Fab constant domain. The system provides a compact test case for particle-specific atomic reconstruction in the presence of flexible regions.

Starting from the full-length sequence of PfCRT and the Fab fragment, we excluded the variable N-terminal (residues 1–46) and C-terminal (residues 406–424) regions of PfCRT that were not modeled in the original structure. Analysis of the latent variable *z* reveals multiple modes of motion in the Fab constant domain (Z1–Z3 in Fig. 3b, left), consistent with regions improved by the heterogeneous refinement (Fig. 3a, right). An additional mode (Z4 in Fig. 3b, right) captures variability in the Fab variable-region loop, likely reflecting local fluctuations in this region; however, this motion is not strongly reinforced by the heterogeneous refinement and presumably corresponds to a lower-amplitude degree of flexibility. Overall, the dominant conformational variability is concentrated in the Fab constant domain, consistent with the missing density in the homogeneous reconstruction (Fig. 3a).

### CryoARC on a benchmark IgG dataset

To compare CryoARC with state-of-the-art continuous heterogeneity methods, we evaluated it on a benchmark dataset from CryoBench [38]. The dataset consists of 100,000 synthetic cryo-EM images of an IgG antibody complex (PDB: 1HZH) generated at a signal-to-noise ratio (SNR) of 0.01. Conformational heterogeneity is introduced by continuously rotating one Fab arm through 360°, producing a one-dimensional circular manifold of motion (Fig. 4a).

**Fig. 4.**
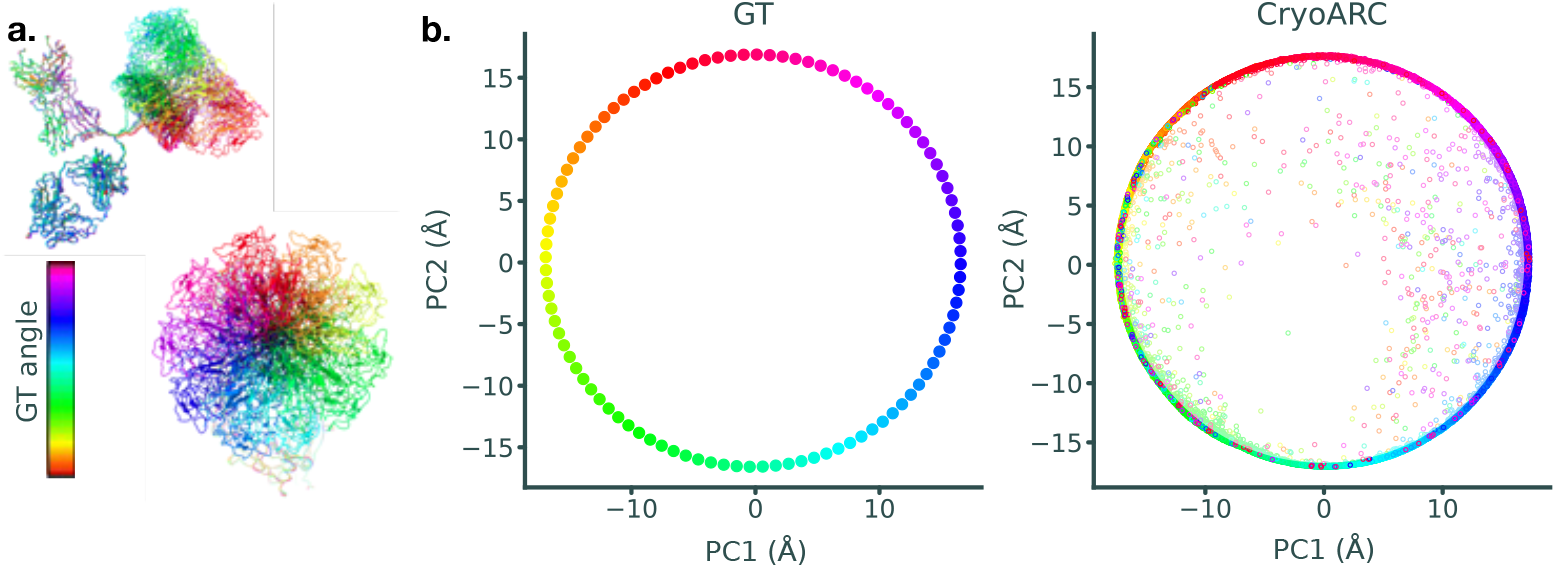
Human immunoglobulin G (IgG) antibody complex dataset from the CryoBench IgG-1D benchmark. **a**. Ground-truth 3D conformational ensemble, colored by the rotation angle of the Fab region, shown in side and top views. **b**. Principal component analysis (PCA) projections of groundtruth conformations (left) and CryoARC-predicted particle-specific conformations (right), shown in a shared embedding space for direct comparison.

We trained CryoARC using the IgG sequence, particle images, and associated poses. To compare predicted and ground-truth conformational spaces, we project particle-level latent variables into atomic conformational coordinates and embed both predicted and ground-truth structures into a shared principal component analysis (PCA) space (Fig. 4b). This procedure enables direct geometric comparison of the recovered conformational manifolds.

CryoARC recovers a conformational space that closely matches the ground-truth circular manifold. The average RMSD between corresponding conformational states is 4.9 Å, which is below the Nyquist limit of the dataset (6 Å). CryoARC achieves comparable recovery of the ground-truth conformational manifold to previously reported methods on this benchmark [38], while additionally producing particle-specific all-atom structural reconstructions rather than volumetric representations.

### Sensitivity of CryoARC to sequence-derived structural priors

We evaluated the sensitivity of CryoARC to sequence-derived structural priors using a synthetic dataset of adenylate kinase (AdK) generated from an all-atom molecular dynamics (MD) simulation of the open–closed transition [39]. We sampled 100 conformational states along the trajectory and generated 100 projection images per state, resulting in 10,000 particles. Images were simulated at 128 × 128 pixel size with a sampling rate of 1.2 Å. The imaging model included a 300 kV electron microscope with a spherical aberration coefficient of 2.7 mm and defocus values uniformly sampled between -0.4 and -0.6 μm.

To assess the influence of sequence-derived structural priors, we repeated the reconstruction using a set of homologous AdK structures with varying sequence identity relative to the ground-truth structure (E. coli AdK, PDB 4AKE). Specifically, we initialized CryoARC using eight homologous structures spanning bacteria, archaea, and eukaryotes: Aquifex aeolicus (PDB 2RH5), Thermus thermophilus (PDB 3CM0), Bacillus subtilis (PDB 1P3J), Mycobacterium tuberculosis (PDB 1P4S), Photobacterium profundum (PDB 4K46), Methanococcus thermolithotrophicus (PDB 1KI9), Saccharomyces cerevisiae (PDB 1AKY), and Notothenia coriiceps (PDB 5X6K). These structures provide a controlled set of sequence-derived priors with varying structural similarity. Figure 5a shows the corresponding homologs and their sequence identities.

**Fig. 5.**
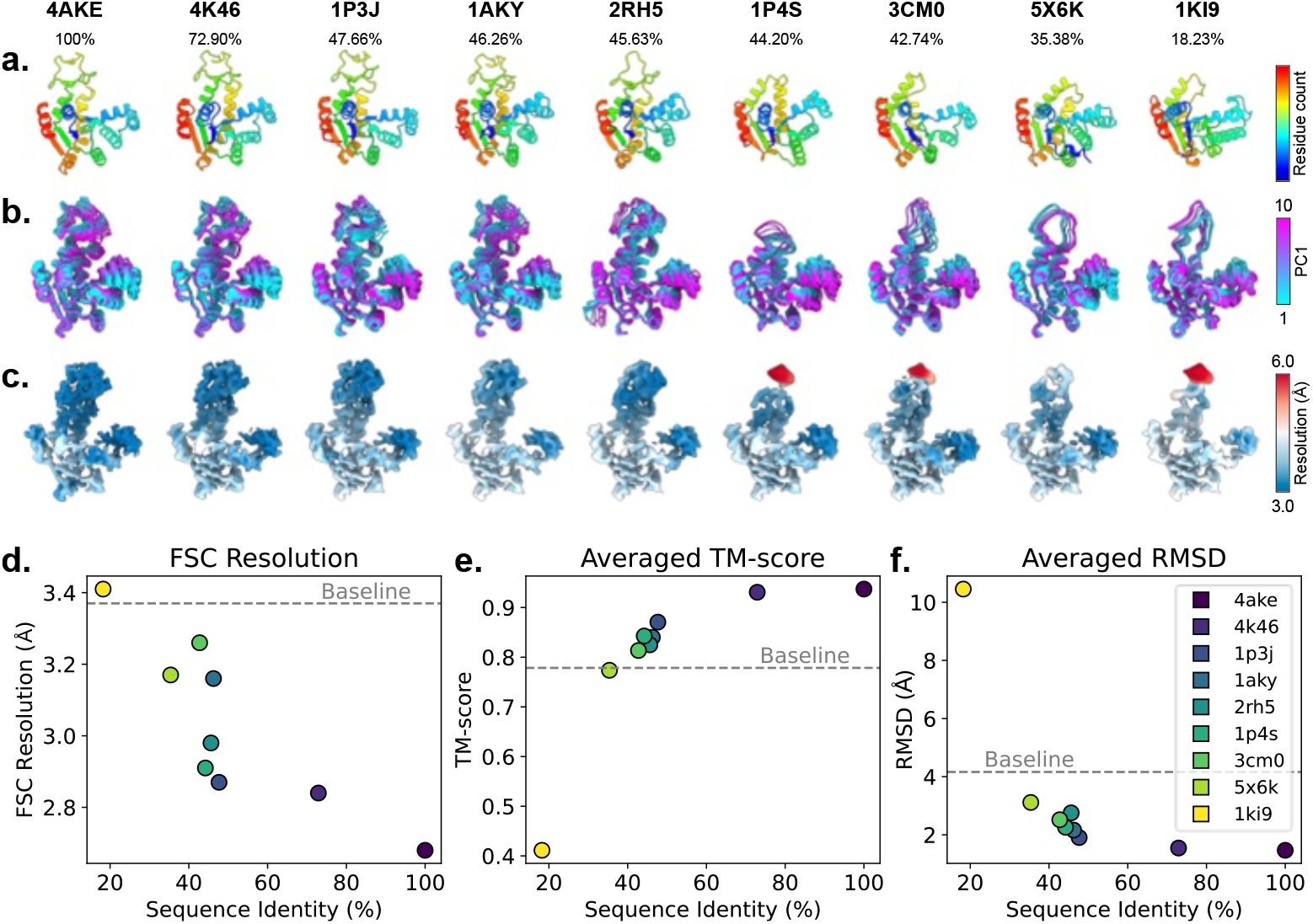
Sensitivity of CryoARC to sequence-derived structural priors. **a**. Homologous AdK structures used as sequence-derived structural priors for CryoARC reconstruction. Structures are ordered by sequence identity relative to the reference AdK structure (PDB 4AKE), from 100% to 18.32%.**b**. Dominant conformational motions recovered by CryoARC along the first latent component for each homologous structural prior. **c**. Local-resolution maps obtained after heterogeneous refinement using CryoARC predictions associated with each homologous prior. **d**. Global resolution estimated from Fourier shell correlation (FSC) after heterogeneous refinement. The baseline corresponds to homogeneous reconstruction without CryoARC-based refinement. **e**. Average TM-score between the ground-truth conformational ensemble and CryoARC-predicted conformations for each homologous prior. The baseline corresponds to the TM-score between the ground-truth ensemble and the static reference structure (PDB 4AKE). **f**. Average RMSD between the ground-truth conformational ensemble and CryoARC-predicted conformations for each homologous prior. The baseline corresponds to the RMSD between the ground-truth ensemble and the static reference structure (PDB 4AKE).

We analyzed the conformational spaces reconstructed under each prior. Across all cases, CryoARC recovered the global open–closed motion of AdK. Regions that are structurally conserved across homologs, such as the LID domain, are consistently well-resolved and show stable conformational trajectories across all initializations. In contrast, regions with variable conservation across the homologs, such as the AMPbd domain, exhibit differences in reconstruction quality depending on the availability of structural information in the prior.

Figure 5b shows the recovered conformational spaces for each homologous initialization. The LID domain motion is consistently captured across all cases. When the AMPbd domain is present in the prior, CryoARC recovers coherent motion in this region. When this domain is absent or structurally divergent, the corresponding region exhibits increased conformational variability and reduced structural consistency in the reconstructed ensemble.

We quantified reconstruction quality using the TM-score and RMSD between the ground-truth conformational ensemble and the reconstructed structures (Fig. 5e,f).Across all initializations, CryoARC achieves improved agreement with the ground-truth ensemble compared to a static reference structure (4AKE). Reconstruction quality correlates with sequence identity between the homologous structural prior and the target structure, indicating that prior–data consistency influences the accuracy of the recovered conformational ensemble.

Finally, we evaluated the effect of structural priors on heterogeneous refinement. Figure 5c shows the resulting density maps for each initialization. Cases with higher sequence identity exhibit higher-resolution refined maps (below 3 Å in Fig. 5d), with well-defined density in both LID and AMPbd domains. In contrast, cases with lower sequence identity show reduced global resolution (above 3 Å) and weaker density in regions not well supported by the structural prior. The LID domain, which is conserved across all homologs, remains consistently resolved with approximately 3 Å local resolution in all cases.

Overall, these results indicate that the quality of the CryoARC reconstruction depends on the coherence between sequence-derived structural priors and the observed cryo-EM data. When structural priors are consistent with the target conformational landscape, CryoARC recovers accurate ensembles and high-resolution density maps. When priors are less consistent, reconstruction remains stable but with reduced structural resolution in less supported regions.

## Discussion

Conventional single-particle cryo-EM heterogeneity analysis typically relies on discrete classification approaches, such as those implemented in Relion [1] and CryoSPARC [2]. While highly effective for resolving distinct structural states, these methods are less suited to describing continuous conformational variability. To model continuous conformational variability, a range of approaches have been proposed, including manifold learning methods [3, 4], covariance- and PCA-based approaches [5, 10, 40], and molecular mechanics–based methods that deform atomic models using normal modes, molecular dynamics, or related representations [6–9].

Recent cryo-EM heterogeneity methods increasingly rely on neural network autoencoders that map particle images to low-dimensional latent representations and decode them into 3D structural models [11–18, 41–43]. In several approaches, the decoder operates directly in voxel or density space, producing 3D reconstructions without explicit structural constraints [11–13]. While this method allows the model to represent a wide range of conformational variability, it can also result in density variations that may not be interpretable as realistic molecular structures. A key challenge, therefore, is to regularize the reconstruction to ensure that the recovered conformational landscapes remain structurally and physically plausible.

Several improvements to autoencoder-based cryo-EM reconstruction have introduced stronger structural regularization. A common strategy is to condition the reconstruction on a reference density map obtained from homogeneous reconstruction, restricting conformational variability to deformations of this volume rather than unconstrained changes in voxel space. For example, 3Dflex represents conformational changes as a deformation field applied to a tetrahedral mesh [41], while methods such as E2GMM and Dynamight model density as a mixture of Gaussian kernels with learned deformations of kernel positions [14, 15]. These approaches introduce geometric constraints that improve stability and interpretability compared to unconstrained voxel-based decoders, but they remain dependent on the quality of the initial density map. In particular, poorly resolved or highly flexible regions in the reference reconstruction can limit the expressiveness of the deformation model and reduce the accuracy of recovered motions. Another extension of autoencoder-based frameworks incorporates structural priors derived from atomic models, obtained either experimentally or predicted in silico [16–18, 42, 43]. These structural priors provide information on local geometry and plausible conformational variations that are difficult to infer from cryo-EM images alone. However, in practice, such constraints are typically limited to simple geometric terms due to the computational cost of explicit energy calculations. As a result, these models must balance fitting accuracy by structural stability: overly unconstrained representations can lead to unrealistic conformations, while overly strong constraints can restrict the accessible conformational space. Recent methods address this trade-off by enforcing local rigidity, for example, through elastic network models or a decomposition of the structure into rigid segments [16–18]. While this improves stability, it can also bias the reconstruction toward the initial structure, limiting the ability to capture larger-scale domain motions. In contrast to previous methods that rely on fixed atomic models or reference densities, CryoARC uses sequence-derived evolutionary embeddings as a structural prior, replacing explicit physics-based constraints (e.g., geometric constraints, force fields, or elastic network models) with an implicit prior learned from evolutionary information. Simple geometric penalties are applied solely to prevent large deviations from configurations suggested by the evolutionary model.

Beyond predicting conformational directions, an additional challenge in heterogeneous cryo-EM analysis is the reconstruction of a single high-resolution canonical map representing the average structure. Several methods address this problem by exploiting predicted conformational variability to improve reconstruction. For example, DynaMight uses particle-specific deformations predicted by the decoder to refine the canonical map [14]. Rather than applying these deformations directly, it trains an additional neural network as a regressor in parallel with the decoder to optimize a least-squares reconstruction objective. Similarly, 3Dflex reconstructs a refined canonical map by applying predicted deformations to a reference density, thereby incorporating particle-specific conformational information into the back-projection step [41]. In CryoARC, we address this problem by incorporating predicted atomic displacements directly into the conventional least-squares reconstruction objective. By explicitly accounting for particle-specific conformational variability during back-projection, CryoARC produces a canonical map that preserves high-resolution features while remaining consistent with the continuous heterogeneity inferred for each particle. The sequence-embedding component in CryoARC is conceptually related to sequence-based protein structure prediction methods, including AlphaFold2 (AF2)

[19] and related models [20–22], which predict atomic structures with high accuracy directly from amino-acid sequences. However, these approaches are primarily designed to generate a single conformation and do not explicitly capture conformational ensembles. To enhance conformational diversity, the community has explored multiple strategies. Some approaches modify the input multiple sequence alignment (MSA) through clustering, subsampling, or mutagenesis to generate alternative conformations [23–26]. Other methods increase the dropout rate in specific AF2 layers, expanding the range of conformations sampled [27]. Generative modeling techniques have also been proposed to emulate dynamics observed in molecular dynamics simulations, including diffusion models [28] and flow matching [29, 30]. Additional strategies leverage structural templates to guide predictions toward specific states [31, 32], or aim to learn protein conformational variability directly from sequence data [33–35]. Several recent methods incorporate experimental data to guide structure prediction toward specific conformational states. Techniques such as cross-linking mass spectrometry [44] and double electron–electron resonance (DEER) [45] provide residue–residue distance constraints that can be encoded into the pairwise representations used by sequence-based predictors, like AF2. Other approaches, such as ModelAngelo, combine multiple sequence alignments with a single cryo-EM density map to produce a static atomic or pseudo-atomic model, mapping residues to the map after an initial model-building step [46]. While these techniques demonstrate that sequence information can improve map interpretation or enforce pairwise constraints, they are limited to either a single map or a single static conformation and do not operate directly on individual particle images. In contrast, CryoARC introduces a novel strategy for incorporating experimental cryo-EM data into generative ensemble predictions by mapping a low-dimensional latent variable learnt from particle images into the pairwise residue representation through the Flexformer network. This allows the model to learn distributions of conformational states that are simultaneously consistent with evolutionary priors and the experimental data, enabling ensemble reconstruction rather than a single static structure.

The main limitation of CryoARC is its computational cost, both in terms of runtime and GPU memory usage. As illustrated in Figure S2, the compute time scales as 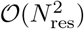, while GPU memory requirements scale as 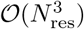.This scaling arises from the use of Invariant Point Attention (IPA) in the decoder, which computes attention weights over all pairs of residues. As a result, the computational cost of CryoARC is substantially higher than that of purely density-based cryo-EM heterogeneity methods. In practice, GPU memory availability constitutes the primary bottleneck. The largest currently available GPUs are limited to approximately 200 GB of memory (e.g., NVIDIA H200), which, according to our scaling analysis, corresponds to a practical upper limit of roughly 3,000 residues. More commonly used GPUs, such as the NVIDIA A100 with 80 GB of memory, restrict the method to proteins of approximately 2,000 residues. Another limitation concerns the initial prediction of multimeric assemblies. In its current form, CryoARC relies on pre-trained Evoformer weights from Alphafold-Multimer to produce structural embeddings. However, AlphaFold-Multimer can exhibit errors in multimer stoichiometry or in predicting inter-chain interfaces, which can strongly influence the downstream reconstruction and degrade the quality of the predicted conformational ensemble. Although CryoARC is capable of modeling conformational variability within multimeric systems, the current framework is not yet robust to changes in assembly state or large inter-chain rearrangements.

Overall, CryoARC introduces a framework that bridges protein structure prediction and cryo-EM heterogeneity reconstruction. While currently limited by computational scaling and applicable to proteins of a few thousand residues, it demonstrates that sequence-informed priors can substantially improve the interpretability and resolution of heterogeneous reconstructions, providing a basis for further integration of experimental data into ensemble structure determination.

## Online Methods

### Sequence encoder

CryoARC embeds the amino-acid sequence of the macromolecular complex by constructing an MSA using a standard database search (see 1). The MSA features are processed through an *Evoformer* module as introduced in AF2 [19]. This module produces residue-wise sequence embeddings 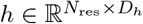 and pairwise residue contact embeddings 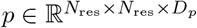, where *N*_res_ is the total number of residues and *D*_*h*_ and *D*_*p*_ are the respective embedding dimensions. It corresponds to Algorithm 6 in [19],where the notations have been adapted, notably (*z, s*) in [19] corresponds to (*p, h*) here.

Importantly, we compute the embeddings *p* and *h* once for each dataset and keep them fixed while training the latent encoder and structure decoder. No gradients are backpropagated through this module, as these embeddings serve purely as structural priors for the latent-conditioned decoder. We obtained the weights in this module from AF2, which was trained to predict static atomic structures as observed in the PDB. In practice, we use parameters from a version of AF2 fine-tuned with templates, without a predicted template modeling score (pTM) from AF2 for single-chain proteins, and from AlphaFold-Multimer v2.3 for protein complexes (see 1).

### Image encoder

To model conformational heterogeneity, we map each particle image *y*_*i*_ to a low-dimensional latent variable *z*_*i*_ ∈ ℝ^*Z*^ describing its conformational state. We use a variational autoencoder (VAE) framework with an encoder distribution *q*(*z*_*i*_| *y*_*i*_) thatapproximates the posterior distribution over latent conformations. The encoder predicts the mean 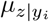 and variance 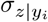 of a Gaussian latent distribution, from which latent samples are drawn using the reparameterization trick:

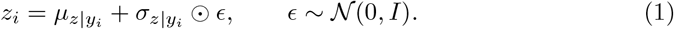

This probabilistic formulation allows a smooth latent representation of conformational variability and captures uncertainty in the inferred states. Particle images are first phase-flipped using the contrast transfer function (CTF) and processed by a convolutional encoder operating in real space. The encoder consists of successive convolution, group normalization, and SiLU activation layers, followed by global average pooling and a linear projection to a 2 × *Z* output representing 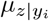 and 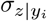.

### Pose encoder

CryoARC jointly refines particle poses and conformations during training. Because pose and conformational variability are strongly coupled in cryo-EM images, ab-initio pose estimation is challenging. We therefore initialize particle poses using conventional cryo-EM alignment procedures (see Supplementary Methods 1), which provide estimates of the viewing directions and in-plane translations for each particle. For each particle image, the pose is represented as *ϕ*_*i*_ = (*θ*_*i*_, *t*_*i*_), where *θ*_*i*_ ∈ℝ^4^ is a unit quaternion encoding the 3D orientation and *t*_*i*_∈ ℝ^2^ is the in-plane translation. The initial pose estimates are denoted 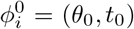. To stabilize training, the particle poses are fixed to 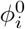 during an initial warm-up phase of *τ*_*ϕ*_ epochs, allowing the latent conformational variables *z*_*i*_ to converge before pose refinement begins. After this stage, the poses *ϕ*_*i*_ are optimized jointly with the latent variables. To prevent large deviations from the initial alignments, pose refinement is regularized with an MSE loss relative to 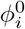.

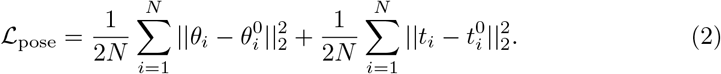

### Latent-conditioned structure decoder

Given the pairwise representation *p* from the sequence encoder and a latent conformation variable *z*_*i*_, the decoder produces an updated pairwise embedding 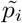 that reflects the conformation encoded in *z*_*i*_. To inject the latent state into the structural representation, we apply a series of cross-attention blocks, which we refer to as the *FlexFormer* module. Each block performs multi-head cross-attention between *z*_*i*_ and the pairwise embedding *p*, first processing along residue rows and then along residue columns, followed by an MLP transition, dropout, and layer normalization:

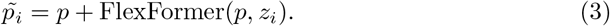

Algorithm 1 provides a schematic representation of the sequence of operations within the FlexFormer module.

#### Algorithm 1

FlexFormer

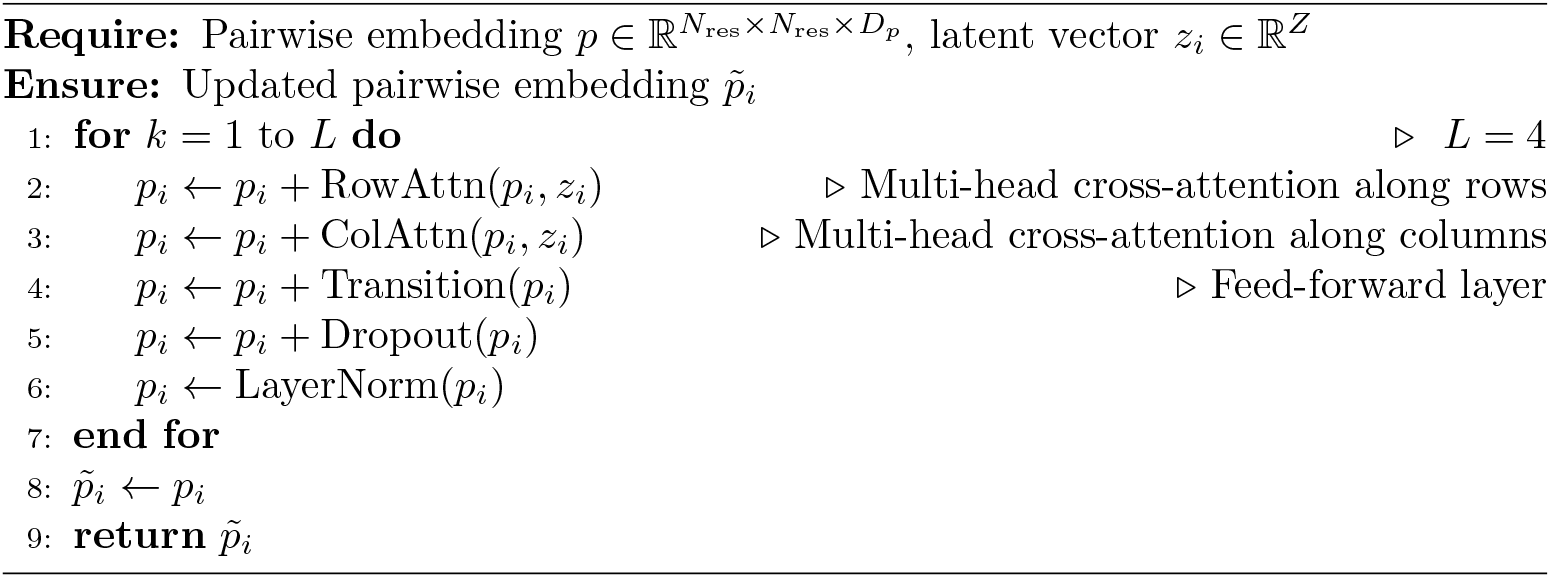

CryoARC then passes the updated pairwise representation 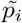 to a pre-trained structure module that predicts the corresponding 3D atomic coordinates,

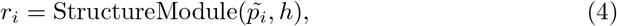

where 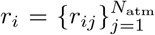 are the predicted atomic coordinates composed of *N*_atm_ atoms with Cartesian coordinates *r*_*ij*_ ∈ ℝ^3^. The structure module closely follows the AF2 architecture, including the IPA block. For training the VAE on heterogeneous cryo-EM particle images, we initialize the IPA parameters using those from AF2. While training, the parameters related to IPA are fine-tuned together with the VAE parameters. During this phase, we keep the side-chain prediction parameters in the structure module frozen to preserve the side-chain predictions of the pre-trained model.

### Cryo-EM forward model

With the predicted atomic coordinates *r*_*i*_ of the *i*-th particle, we can simulate the cryo-EM image formation to obtain a projected image *x*_*i*_ that we will use for loss computation. Cryo-EM particle images measure the 3D Coulomb potential of the sample projected along the electron beam direction, modulated by the microscope’s contrast transfer function (CTF). Let 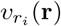 denote the discrete 3D Coulomb potential resulting from the atomic structure with coordinates *r*_*i*_ at voxel position **r** and 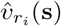 its continuous Fourier transform at spatial frequency **s**. Given the pose parameters *ϕ*_*i*_ = (*θ*_*i*_, *t*_*i*_) where *θ*_*i*_ *SO*(3) is the particle orientation and *t*_*i*_ ∈ ℝ^2^ is an in-plane 2D translation, the noise-free Fourier component of the projected image at the in-plane 2D spatial frequency **s** = (*s*_*x*_, *s*_*y*_, 0)^⊤^ is

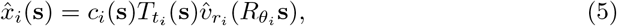

where *c*_*i*_(**s**) denotes the CTF, 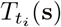 is a phase shift corresponding to a 2D translation of *t*_*i*_ in real space, and 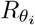is the 3D rotation corresponding to the particle orientation *θ*_*i*_. Here, 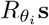 produces a point in the 3D Fourier volume along the central slice oriented by *θ*_*i*_.

To simulate potential 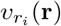from the atomic structure with coordinates 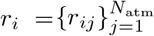 composed of *N*_atm_ atoms with Cartesian coordinates *r*_*ij*_ ∈ ℝ^3^, we approximate the Coulomb potential as a superposition of isotropic Gaussian kernels centered at the atomic coordinates,

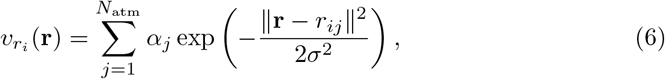

where *α*_*j*_ scales the Gaussian kernels to model the electron scattering strength of the *j*-th atom, and *σ* controls the width of the Gaussian kernels and models the spatial blurring. We set *α*_*j*_ to the atomic number of the corresponding atom, which roughly follows the number of electrons in the atom, to simulate its scattering strength. We selected *σ* = 1 Åso that the simulated density roughly reflects the blurring of the electron microscope at high resolution. Using properties of Gaussian functions and their Fourier transform, we can express the 3D density in the Fourier domain as

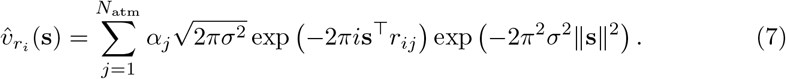

### Training

CryoARC is trained within a variational autoencoder (VAE) framework. For each particle image *y*_*i*_, the encoder predicts a latent distribution *q*(*z*_*i*_ | *y*_*i*_), and the model is optimized using a reconstruction objective together with a Kullback–Leibler regularization term:

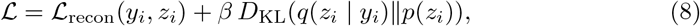

where *p*(*z*_*i*_) = 𝒩 (0,*I*) is a standard Gaussian prior over latent variables. The expectation over *q*(*z*_*i*_ | *y*_*i*_) is approximated using a single Monte Carlo sample with the reparameterization trick (Eq. 1). Instead of using a standard pixel-wise Gaussian reconstruction loss, which is sensitive to the global scaling of the image intensities, we avoid normalizing the images and rather optimize a normalized correlation objective in Fourier space:

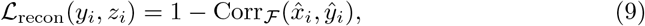

where

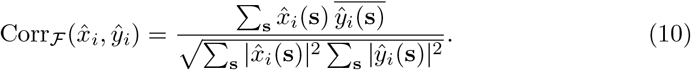

Here, 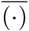 denotes complex conjugation and 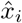and 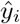 denote the Fourier transforms of the predicted and experimental images, respectively. This loss is equivalent to a complex-domain cosine similarity and is invariant to the global scaling of the image intensities while remaining sensitive to their phase differences.

In addition to the data reconstruction loss, we utilize two structure preservation losses, a side chain and backbone torsion angle loss ℒ_Torsions_, and a structural violations loss ℒ_Violations_ as defined in AF2 [19]. The ℒ_Torsions_ loss penalizes differences in torsion angles between the predicted structures and the ground truth. Since we assume that the ground-truth structure is unknown during reconstruction, we use this loss to compare the geometry of the structure predicted with the latent conditioning against the structure predicted without it. This loss helps ensure that the model parameters remain close to those of the initial pre-trained structure-prediction model. The ℒ_Violations_ loss penalizes the bond lengths, bond angles, and atomic clashes against reference values from crystallographic tables [47]. The total auxiliary loss reads as

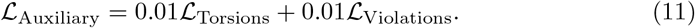

We train the parameters of the Encoder, FlexFormer, and IPA end-to-end using the Adam optimizer. Details regarding the training parameters (e.g., learning rate, number of epochs, and number of parameters) are available in the Supplementary Information.

### Heterogeneous reconstruction

To reconstruct a canonical 3D density from particle images with heterogeneous conformations, we introduce a deformation-aware backprojection, guided by the predicted per-particle atomic positions. It maps the contribution of each atom in each particle into a single canonical volume. In short, this procedure deforms each back-projected image according to its predicted atomic coordinates into the coordinates of a canonical structure. To account for the CTF modulation and noise associated with back-projection, we redefine the Wiener filtering reconstruction, commonly used in cryo-EM.

We begin by defining a canonical molecular structure with atom coordinates 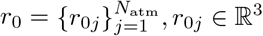, obtained by decoding the average latent variable 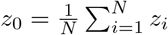. We note Δ*r*_*i*_ = *r*_*i*_ − *r*_0_ the per-particle atomic displacement associated with the *i*-th particle. Let *v*_0_ be the unknown density of the canonical volume associated with the canonical coordinates *r*_0_ that we aim to reconstruct. Given Δ*r*_*i*_, we define an interpolation operation *g*(*v*, Δ*r*_*i*_), that for a given particle *i*, shifts and interpolates the per-atom density:

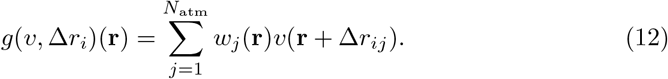

Here, *w*_*j*_(**r**) is a Gaussian interpolation that acts as a narrow window centered around the *j*-th atom, defined as

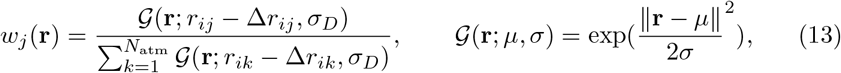

where 𝒢 is a 3D Gaussian kernel, and *σ*_*D*_ controls the smoothing of the interpolation. The interpolation function *g* effectively focuses the density locally around the shifted density *v*(**r** + Δ*r*_*ij*_); however, it does not account for the density distant from the atomic coordinates, such as background or unmodeled atoms (e.g., ligands). Therefore, to preserve the background signal in density regions distant from the predicted atomic coordinates, we introduce a masking strategy defined below (dependency on position has been omitted for readability)

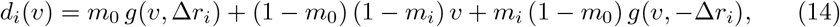

where *m*_*i*_ and *m*_0_ are masks that set the regions inside the molecule to unity and those outside to zero, defined as

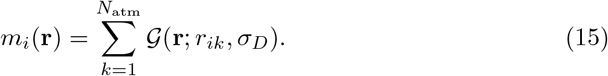

The effect of this masking strategy is visually described in Figure S3. Next, we approximate the interpolation function *d*_*i*_(*v*)(**r**) as being diagonal in the Fourier space,

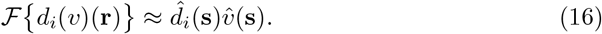

Note that since *w*_*j*_ and *m* vary with **r**, the Fourier transform in Eq. 16 involves convolutions with 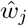 and 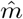, implying that *d*_*i*_ is generally not strictly diagonal in the Fourier domain.

Using the newly introduced interpolation function *d*_*i*_(*v*)(**r**), we can write down the formation model defined in Eq. 5 as a function of the canonical density volume 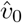 as

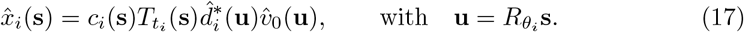

Note that we use the complex conjugate of the Fourier transform of *d*_*i*_(*v*)(**r**) in the forward model. Indeed, since we defined the interpolation as the inverse mapping from the deformed particle back to the canonical density, in the Fourier domain, it corresponds to the Hermitian transpose.

With the forward model defined above, we can find an estimate of the minimum mean squared error (MMSE) problem, also known as Wiener filtering [1, 48, 49]. We assume that the normalized observed images are corrupted by an additive noise, with zero-mean and unit variance uncorrelated between frequencies 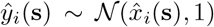, and we also assume a Gaussian prior on the canonical volume 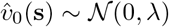where we approximate the power spectrum to be a constant *λ*, as commonly accepted in the cryo-EM models [50]. With the assumptions above, we can write the MMSE problem as

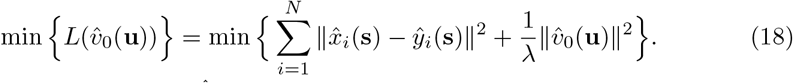

By defining 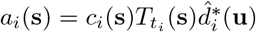 and substituting Eq. 17, we can write

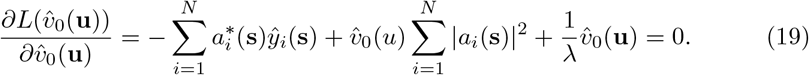

The solution 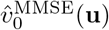has the common form,

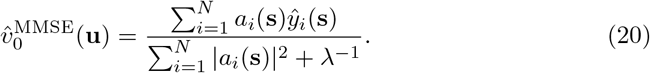

By substituting *a*_*i*_(**s**) into the above equation, we obtain

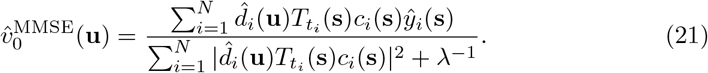

Finally, we calculate the interpolation in real space. Using Eq. 16, we can write the final form of the MMSE solution,

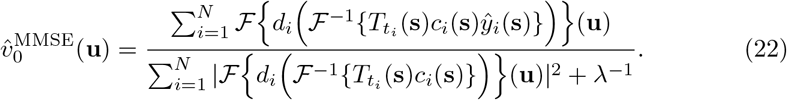

### Structure initialization

In addition to sequence-derived evolutionary embeddings, CryoARC can optionally be initialized using an initial atomic structure when sequence information alone is insufficient to provide a reliable prior. This setup is particularly relevant in multimeric systems, where intra-chain geometry is often well predicted, but inter-chain orientations and interfaces can be inaccurate.

When a structural prior is provided, we first pre-train the structure decoder without conditioning on the latent conformational variable *z*_*i*_. During this stage, the model is trained to reproduce the input structure using a frame-aligned point error (FAPE), following AF2 formulations [19]. This pre-training is performed for a small number of epochs to initialize the decoder in a physically reasonable region of conformational space.

After this pre-training phase, latent conditioning is enabled and full CryoARC training proceeds. This procedure is used in all experiments that feature multimeric systems.

### Evaluation metrics

We calculated local resolution densities with CryoSPARC [2] with a threshold of 0.5 RMSD values were calculated between C_*α*_ atoms of ground-truth and predicted structures and averaged over the entire dataset. TM-scores were calculated using the tools available from the Zhang lab at https://aideepmed.com/TM-score/.

## Acknowledgments

This study was supported by the French National Research Agency (research grant ANR-23-CE45-0024-01) and within the framework of the “Investissements d’avenir” program (ANR-15-IDEX-02).

## Supplementary Methods

### Genetic search and Multiple Sequence Alignment (MSA)

We generate sequence embeddings following the pipeline described in the AF2 publication [19], with the OpenFold implementation [51] used in inference mode. Starting from the protein sequence in FASTA format, Openfold searches multiple genetic databases with JackHMMER [52] and HHblits [53], and constructs the MSA of homologous sequences. We filter, crop, and featurize the resulting MSA following the procedure described in the AF2 publication [19], and then embed it in the Evoformer block.

### Sequence encoder and structure decoder parameters

The parameters of the sequence encoder and structure decoder in CryoARC are exactly the parameters of the *Evoformer* and *Structure Module* in AF2 [19], respectively. For monomeric proteins, we used parameter weights obtained from Openfold training, after finetuning, without templates and without predicted Template Modeling (pTM) scores (referred to as *finetuning no templ 1*). For multimers, we used parameter weights obtained from AlphaFold-Multimer v2.3 [54] without templates and without pTM (referred to as *model 3 multimer v3*).

### Pre-alignement of particle stacks and CTF estimation

In cryoARC, we assume that an initial pre-alignment has been performed on the particle stacks (i.e., 3D reconstruction), resulting in particle pose parameters (orientation and shifts) for each particle. We also assume that the CTF parameters for each particle have been pre-estimated.

In the experiments presented above, we used the particle pose and CTF parameters obtained with different cryo-EM single-particle analysis pipelines. For the HER2 dataset (EMPIAR-11665 [36]), the original study utilized CryoSPARC [2] for CTF estimation and 3D reconstruction. For the PfCRT dataset [37] (EMPIAR-10330), the original study utilized CTFFind4 [55], GCTF [56], and cisTEM [57] for the CTF estimation and CryoSPARC and Relion [1] for 3D reconstruction. For the synthetic IgG dataset [38], we used ground-truth CTF parameters and the particle pose provided by the authors of the benchmark. For the AdK dataset, we obtained the particle pose parameters by applying *ab initio* reconstruction and homogeneous refinement functions from CryoSPARC v4 to the particle stacks. We utilized the CTF parameters identical to those used to generate the images (i.e., the ground-truth parameters).

### Training details

We optimized several CryoARC training parameters for each of the datasets presented above. Table 1 outlines this information, including data about the particles, model architectures, and training hyperparameters.

## Supplementary Figures

**Fig. S1.**
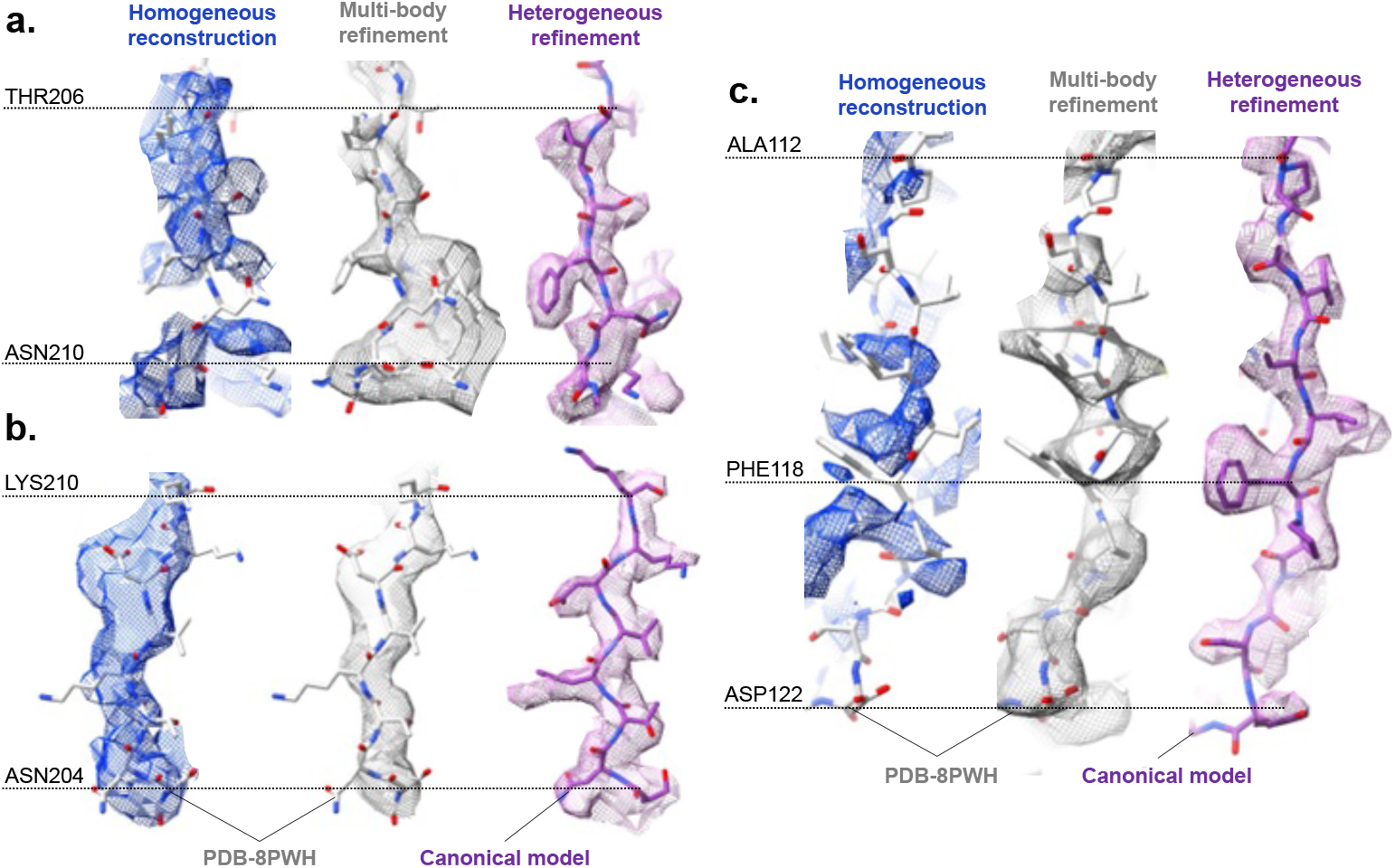
Structural details of the heterogeneous reconstruction on the HER2 dataset in selected highly flexible regions. Blue densities correspond to the classic homogeneous reconstruction (EMD-18188), white densities correspond to a composite map obtained using multibody classification (EMD-17993), and pink densities correspond to CryoARC heterogeneous reconstruction. The overlapped atomic models are PDB-8PWH (grey) and the canonical model from CryoARC. **a**. Residue range THR206 –– ASN210 in chain A. **b**. Residue range ASN204 –– LYS10 in chain B. **c**. Residue range ALA112 –– ASP122 in chain C.

**Fig. S2.**
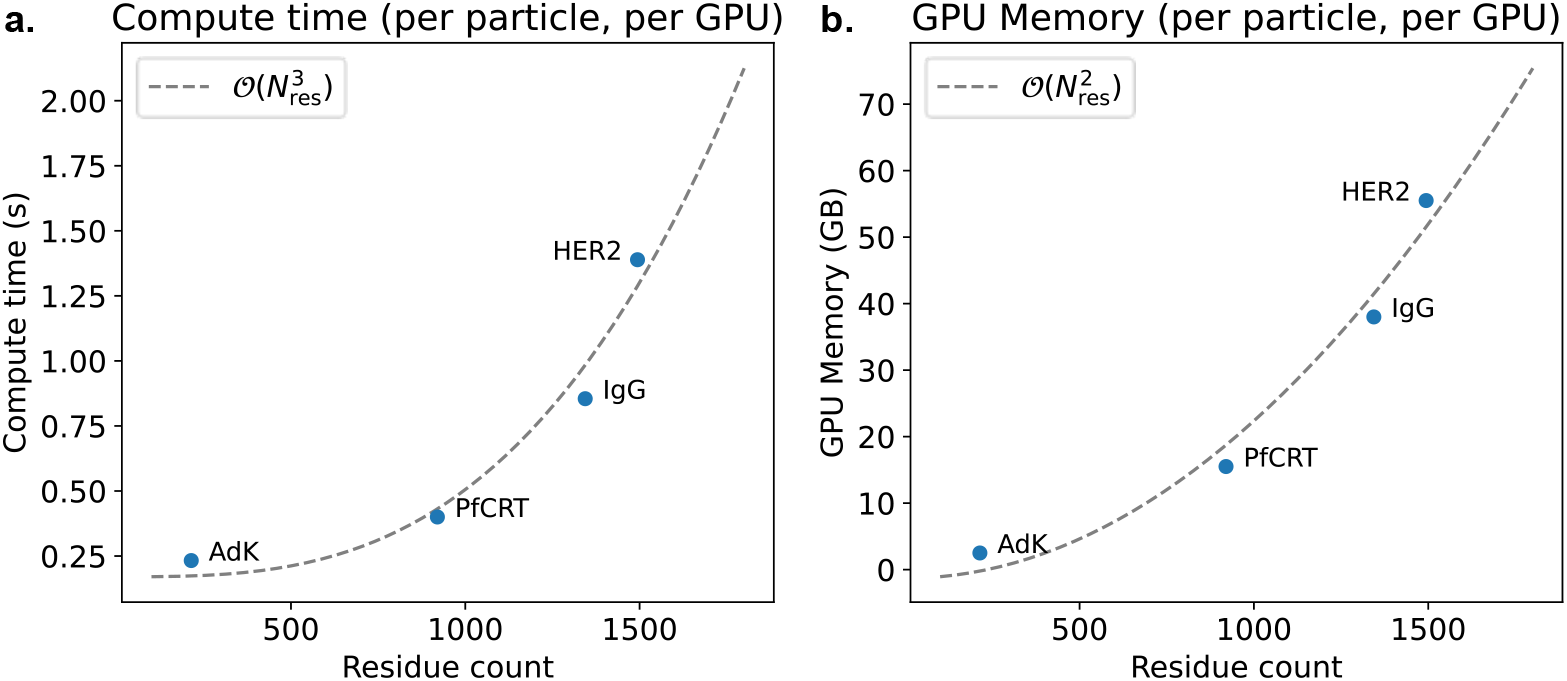
CryoARC computational complexity measured on four datasets, AdK, PfCRT, IgG, and HER2. Dashed lines show polynomial fits. **a**. Compute time as a function of residue count. **b**. GPU memory as a function of residue count.

**Table S1.** Particles, architecture and training details of CryoARC on each of the presented datasets.

| Dataset | IgG | HER2 | PfCRT | AdK |
| --- | --- | --- | --- | --- |
| **Particles** |  |  |  |  |
| Number of particles | 100,000 | 708,610 | 16,904 | 10,000 |
| Particle dimension | 128 | 280 | 220 | 128 |
| Particle sampling rate (Å/pix) | 3.0 | 1.16 | 1.03 | 1.0 |
| **Architecture** |  |  |  |  |
| Encoder layers / hidden dimension | 5/32 | 6/24 | 6/24 | 5/24 |
| Latent dimension | 4 | 4 | 4 | 4 |
| FlexFormer layers / hidden dimension | 4/128 | 4/128 | 4/128 | 4/128 |
| Structure decoder layers / hidden dimension | 4/128 | 4/128 | 4/128 | 4/128 |
| **Training** |  |  |  |  |
| Epochs | 15 | 3 | 100 | 100 |
| Steps (epochs × particles) (million steps) | 1.5 | 2.1 | 1.7 | 1.0 |
| Learning rate | $8 \cdot 10^{-5}$ | $8 \cdot 10^{-5}$ | $5 \cdot 10^{-5}$ | $5 \cdot 10^{-5}$ |
| Pose learning rate | $10^{-4}$ | $10^{-4}$ | $10^{-4}$ | $10^{-4}$ |
| Number of H100 GPUs | 16 | 16 | 12 | 12 |
| Mini-batch size | 32 | 16 | 48 | 256 |

**Fig. S3.**
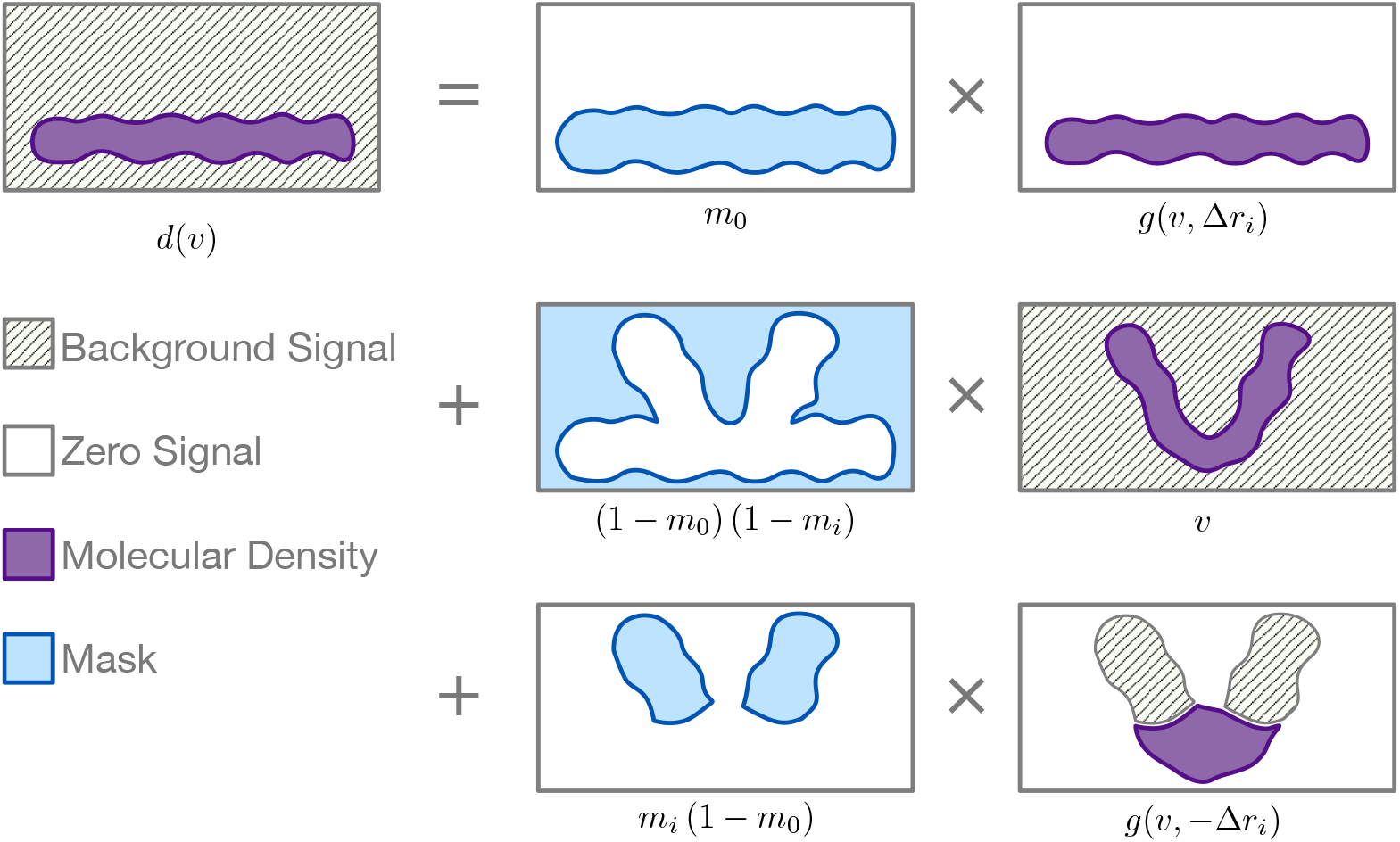
The visual representation of the masking strategy in heterogeneous backprojection illustrates how we combine contributions from molecular density (first row) with the background (second and third rows). This approach enables us to reconstruct the signal beyond the range of the modeled atoms.

